# Allograft and Autograft Anterior Cruciate Ligament Reconstructions Exhibit a Similar Biological Response to Cyclic Loading

**DOI:** 10.1101/2024.03.20.585695

**Authors:** Lauren Paschall, Ariane Tsai, Erdem Tabdanov, Aman Dhawan, Spencer Szczesny

## Abstract

**Objective:** Anterior cruciate ligament (ACL) reconstruction is one of the most commonly performed orthopaedic procedures. While outcomes are similar in the general patient population, the rerupture rate of non-irradiated allografts are 3-4 times higher than autografts in young active individuals. Previous studies suggest that the difference in clinical performance between graft types is due to impaired remodeling in allografts in response to loading. The objective of this study was to compare the remodeling response of autografts and allografts to cyclic loading. Furthermore, given that allografts are a foreign object and that immune cell signaling affects fibroblast mechanobiology, we compared markers of the immune cell composition between graft types.

**Methods:** ACL reconstructions were performed on New Zealand white rabbits, harvested 8 weeks post-surgery, and cyclically loaded to 2 MPa in a tensile bioreactor. Expression of markers for anabolic and catabolic tissue remodeling, as well as inflammatory cytokines and immune cells, were quantified using quantitative reverse transcription polymerase chain reaction.

**Results:** We found that the expression of markers for tissue remodeling were not different between allografts and autografts. Similarly, we found that the expression of markers for immune cells were not different between allografts and autografts.

**Conclusions:** These data suggest that the poor clinical outcomes and impaired remodeling of allograft reconstructions compared to autografts is not due to a difference in graft mechanobiology.

## Introduction

Anterior cruciate ligaments (ACLs) are one of the most commonly torn ligaments affecting more than 200,000 people in the United States each year^1,2^. Given the lack of spontaneous tissue repair and to avoid risk of secondary knee damage^3,4,5^, reconstruction of the ACL is the gold-standard following rupture. Graft options for reconstructing the ACL come in the form of autogenous or allogenic tendons, which have similar outcomes across the general patient population^6,7,8^. However, in young active individuals, who account for nearly 70% of all ACL reconstructions^9,10^, the rerupture rate of non-irradiated allografts is 3-4 times higher than autografts^11,12,13^. This heightened rerupture rate in allografts is likely not due to differences in graft tissue properties since, prior to surgery, the mechanics of non-irradiated allografts and autografts are identical^14,15,16,17^. However, shortly after implantation, the mechanical properties of both graft types dramatically decreases as they undergo significant structural remodeling in a process termed “ligamentization”^18,19,20,21,22^. Over time, the grafts regain their mechanical properties, but this mechanical recovery is slower in allografts compared to autografts^23,24,25^. This suggests that impaired remodeling is the cause of increased allograft rerupture and their poor clinical performance.

One possible cause of impaired allograft remodeling is that allografts induce a deficit cellular response to mechanical loading. There is clear evidence that the remodeling of ACL reconstructions is sensitive to mechanical loading^26,27,28^, and animal studies suggest that allografts and autografts respond differently to mechanical stimulation^23^. Additionally, while clinical data show that rerupture rates are increased in highly active patients for both graft types, the risk ratio with patient activity (and presumably greater mechanical loading) is 14.10 times higher for allograft than autografts^29^. These data suggest that cells within autografts have an impaired response to mechanical loading than those in allografts. However, the reason for this difference in cellular response is not immediately obvious since the cells in both graft types have the same origin. Multiple animal studies demonstrate that, very shortly following surgery, the native cells in autografts die and are replaced by host cells infiltrating the graft tissue^19,21,30^. Still, given that allografts are a foreign object, it is likely that the specific composition of the infiltrated cells (and specifically the immune cells) may be different^24,31^. Interestingly, macrophages and T cells influence the mechanobiology and differentiation of fibroblasts into myofibroblasts, ^32^ suggesting that differential immune cell composition may impair the allograft remodeling in response to mechanical loading. Consistent with this, previous studies demonstrate that allografts have a decreased myofibroblast cell population compared to autografts^23,33^. Since myofibroblasts are responsible for remodeling fibrous tissue^34^ and mechanical loading is needed to induce myofibroblast differentiation^35^, these data suggest that impaired allograft remodeling is due to an impaired cellular response to mechanical stimuli.

Therefore, the objective of this study was to compare the cellular response of autografts and allografts to cyclic loading. We hypothesized that allografts exhibit an impaired remodeling response to cyclic loading (i.e., increased catabolic and inflammatory gene expression) compared to autografts. Additionally, we also hypothesized that allografts contain elevated levels of inflammatory immune cells (e.g., M1 macrophages), which may explain the impaired mechanobiological response of allografts. To test these hypotheses, we utilized a previously published explant model^36^ to cyclically load autograft and allograft explants and quantify gene expression between autografts and allografts in response to mechanical loading.

## Materials and Methods

### ACL Reconstruction

Using an established surgical model^37^, 26 male New Zealand white rabbits (2.8–3.2 kg and 14–16 weeks old) underwent unilateral ACL reconstructions (14 autograft and 12 allograft) using the semitendinosus tendons from an IACUC approved study. Briefly, the native ACL was resected, and drill tunnels were created through the femur and tibia at the ACL insertions. The graft was advanced through the tunnels, and the ends of the graft were sutured under slight tension to the periosteum with the knee in full flexion. In animals assigned to autograft reconstructions, the ipsilateral semitendinosus tendon was used. For animals assigned to allograft reconstructions, the semitendinosus tendon was harvested, flash frozen in liquid nitrogen, and stored at −80 °C, while tendons from a donor rabbit were thawed and used for the reconstructions. Rabbits were euthanized and grafts were harvested 8 weeks post-reconstruction. This timepoint was chosen because it marks the end of the initial remodeling phase involving host cell infiltration and revascularization in rabbits^19,21,30^, and it represents a transition point in the remodeling process where the grafts begin to recover their mechanical properties^38,39,40^.

### Cellularity

Two autograft reconstructions were used to confirm that the 8-week timepoint following surgery was sufficient time to allow for host cell infiltration. Following harvest, the reconstructions were sharply dissected from the femur and tibia, fixed with 4% paraformaldehyde, and embedded in paraffin. The entire reconstruction was sectioned (10 mm thickness), stained with DAPI, and coverslipped. Images (509 µm x 383 µm) were acquired at three locations along the sample length (femur side, middle, tibia side) throughout the entire depth of the reconstruction and imaged using an inverted fluorescence microscope (Keyence BZ-9000E). The midusbtance was defined as the sections that were obtained halfway through the reconstruction (approximately 700 mm deep) and the surface was defined as a few sections into the reconstruction (approximately 70 mm deep). Cell density was calculated by counting the number of cells using ImageJ and dividing by the image area. The contralateral native ACLs from each rabbit were used as control samples for comparison of native cell density.

### Mechanical Loading

Following harvest, 12 autografts and 12 allografts were prepared for mechanical testing under sterile conditions following a previously published protocol^36^. Briefly, all surrounding tissues were removed, and the femur and tibia were cut down into bone blocks to grip the sample during mechanical loading. The initial length and major and minor diameters were measured using calipers. The cross-sectional area was calculated by assuming an elliptical cross-section. The reconstructions were placed in a custom tensile bioreactor^41^ with culture media (low-glucose DMEM, 2% penicillin-streptomycin, 5% FBS, 25 mM HEPES, 4mM GlutaMAX, 1 mM 2-Phospho-L-ascorbic acid trisodium) and kept at 37°C and 5% CO_2_. Since underloading is detrimental to tendon/ligament homeostasis^42,43^, a 0.1 MPa static load was applied for 18 hours to acclimate the reconstructions to culture conditions^36^. The bioreactor then cyclically loaded the samples to 2 MPa at 0.5 Hz for 8 h. Control samples were maintained under a 0.1 MPa static load for the same duration. The duration and frequency of loading were chosen based on prior studies of native ACL explants^36^. The 2 MPa stress level was chosen since it is representative of physiological in situ ACL loading and since preliminary testing found that it was the maximum stress that the reconstructions could sustain for 8 h of cyclic loading.^36^

### Macroscale Tissue Strain

Macroscale tissue strain was calculated using linear variable differential transformers (LVDTs) built into the bioreactor as previously described^41,44^. Briefly, tissue displacement was measured by the LVDTs attached to each sample grip, and the macroscale tissue strain was calculated by dividing the peak displacement of each sample at the end of the 8-hour loading period by its initial length. To determine how much of the macroscale strain was due to tissue creep/slippage at bone interface, the creep of each sample was calculated as (Eq. 1):

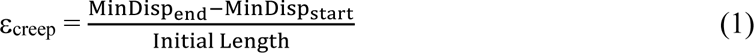

where MinDisp_end_ and MinDisp_start_ are the minimum displacement at zero load during the final and the first cycle, respectively.

### Gene Expression Analysis

After loading at each condition, samples (n = 6) were immediately removed from the bioreactor, and the reconstructions were sharply dissected from the bone blocks. The reconstructions were rinsed with ice cold RNase-free water and then flash frozen in liquid nitrogen. The tissue was pulverized (6775 Freezer/Mill SPEX) and the total RNA was extracted with RNeasy minicolumns (RNeasy Fibrous Tissue Kit, Qiagen). RNA concentration was quantified using a Qubit 4 Fluorometer (Thermo Fisher), and RNA integrity numbers (RINs) were determined using a TapeStation (Agilent) (average RIN of 7.5). Total RNA was diluted to 1 ng/ul, and cDNA was synthesized with a commercially available kit (High-Capacity cDNA Reverse Transcription Kit with RNase Inhibitor, Thermo Fisher).

Quantitative PCR (qPCR) was performed using TaqMan probes (**Supplementary Table 1**) and a StepOne Plus Real Time PCR system to measure the expression of anabolic (C*OL1A1, COL1A2, LOX, COL3A1, TGFβ1 ACTA, TIMP1, TIMP3),* catabolic (*MMP1, MMP2, MMP10, MMP13*), and inflammatory (*IL-1β, PTGS2)* genes. Additionally, markers of M1 macrophages (*NOS2*), M2 macrophages (*MRC1*), helper T cells (*CD4*), and neutrophils (*CXCR2*) were measured along with *GAPDH* as the reference gene. PCR efficiency and cycle number quantification (Cq) were obtained for each individual reaction using PCR-Miner (version 4.0)^45^. Samples that exhibited undetectable fluorescence for a given probe within 50 cycles were excluded from the analysis for that gene. An outlier test (ROUT) was conducted to determine if any of the reaction efficiencies for each gene was an outlier. After removal of outliers, a single efficiency value was calculated for each probe by averaging all the individual sample efficiencies for each gene. Gene expression was quantified using the delta-delta Cq method correcting for primer efficiencies^46^. The fold change for each sample was calculated as (Eq. 2):

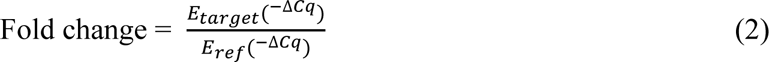

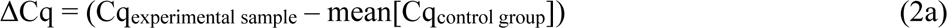

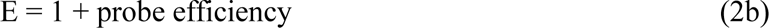

where the probe efficiencies are listed in (**Supplementary Table 1**). To determine the effect of mechanical loading on reconstruction type, the autograft and allograft cyclically loaded samples were compared to their graft-matched statically loaded control samples. To determine baseline graft-specific gene expression differences, the allograft static control samples were compared to the autograft static control samples. To determine baseline differences between reconstructions and the native ACL, allograft and autograft static control samples were compared to previous data acquired from native male ACL static control samples^36^.

### Statistical Analysis

The fold change data for each gene was normalized using a log transformation, and Mann-Whitney tests were conducted on the normalized data to compare the gene expression between graft types. Additionally, Mann-Whitney tests were used on the transformed data to determine the differential expression of each gene compared to the respective control condition (i.e., static load). F-tests were also used to determine whether the reconstructions exhibited greater noise/variability of their gene expression at baseline (i.e., static control) compared to native ACLs. Given the numerous genes evaluated via these statistical comparisons, all p-values were corrected using a false discovery rate. Specifically, to make conclusions about each form of remodeling and the immune cell response, the multiple statistical comparisons conducted within each gene category (anabolic, catabolic, inflammatory, and immune cells) were corrected with a Q-value of 5%. Mann-Whitney tests were used to determine if there was a difference in cross-sectional area, length, macroscale strain, and creep strain between allografts and autografts. Statistical significance after corrections was set as p < 0.05 with statistical trends set at p < 0.10. All statistical analysis was performed using GraphPad Prism (version 9.3.1).

## Results

### Cellularity

After 8 weeks post-reconstruction, the cellularity in the autograft reconstructions (surface: 2848 ± 79 cells/mm^2^, midsubstance: 2172 ± 437 cells/mm^2^) was similar to that of the contralateral ACLs (surface: 2658 ± 1212 cells/mm^2^, midsubstance: 2272 ± 504 cells/mm^2^) (**Fig. 1**).

**Figure 1:**
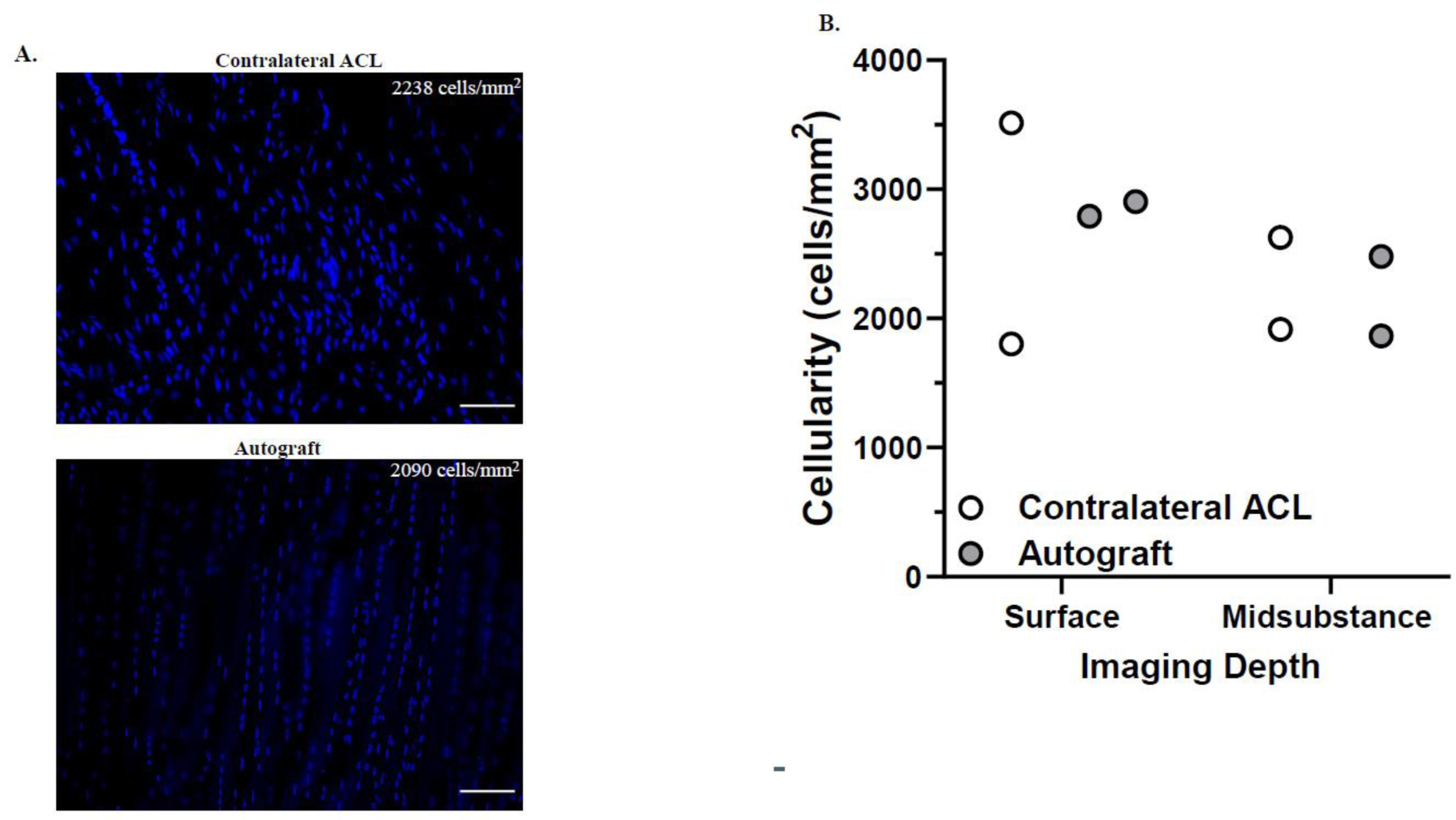
Cellularity of autograft reconstructions 8 weeks post-op. (A) Representative images in the surface and midsubstance of the tissue for autografts and the contralateral ACL. Blue (DAPI) represents cell nuclei (scale bar = 60 µm). (B) Quantification of cell viability for each condition in the surface and midsubstance of the tissue (n = 2 for autograft and contralateral ACL). Data represented by dot plots.

### Remodeling response of ACL reconstructions to cyclic load

At baseline in the statically loaded samples, there was no differential expression of any of the anabolic, catabolic, or inflammatory genes between graft types (**Fig. 2**). Following cyclic loading to 2 MPa, neither autograft nor allograft reconstructions exhibited differential gene expression compared to static controls (**Fig. 3**). Direct comparison between autografts and allografts found that there was no difference in the effect of cyclic loading on gene expression between graft types (**Fig. 4**).

**Figure 2:**
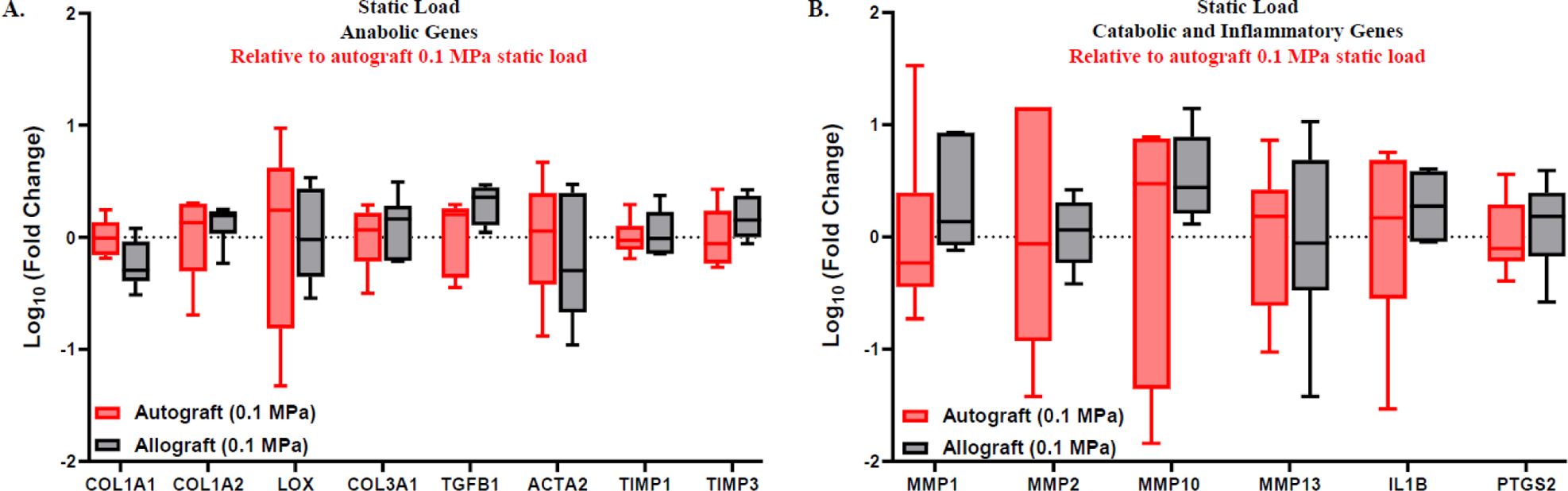
Comparison of autograft and allograft reconstructions gene expression at baseline. RT-qPCR analysis of statically loaded allograft reconstructions relative to statically loaded autograft reconstructions (represented by the red box and whisker plots). (A) Gene expression of anabolic markers. (B) Gene expression of catabolic and inflammatory markers. (n = 6). Data represented as box and whiskers plot with the whiskers representing the min and max data.

**Figure 3:**
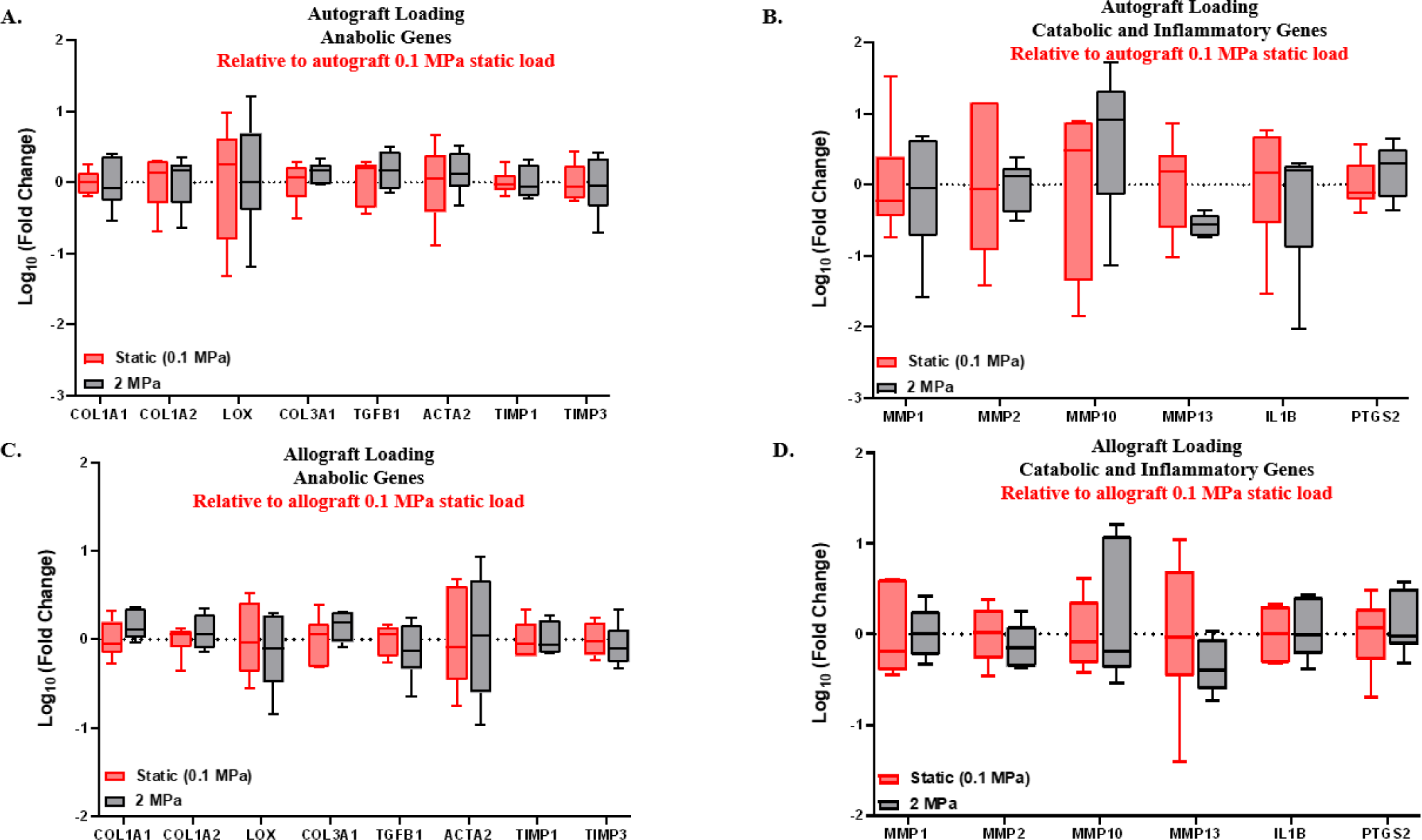
Gene expression changes in ACL reconstructions with mechanical loading. RT-qPCR analysis of autograft and allograft reconstructions after being cyclically loaded relative to the graft-matched 0.1 MPa static load (represented by the red box and whisker plots). (A) Gene expression of anabolic markers and (B) catabolic and inflammatory markers for cyclically loaded autograft reconstructions relative to statically loaded autografts. (C) Gene expression of anabolic markers and (D) catabolic and inflammatory markers for cyclically loaded allograft reconstructions relative to statically loaded allografts. (n = 5-6 for cyclically loaded autografts and n = 6 for cyclically loaded allografts). Data represented as box and whiskers plot with the whiskers representing the min and max data.

**Figure 4:**
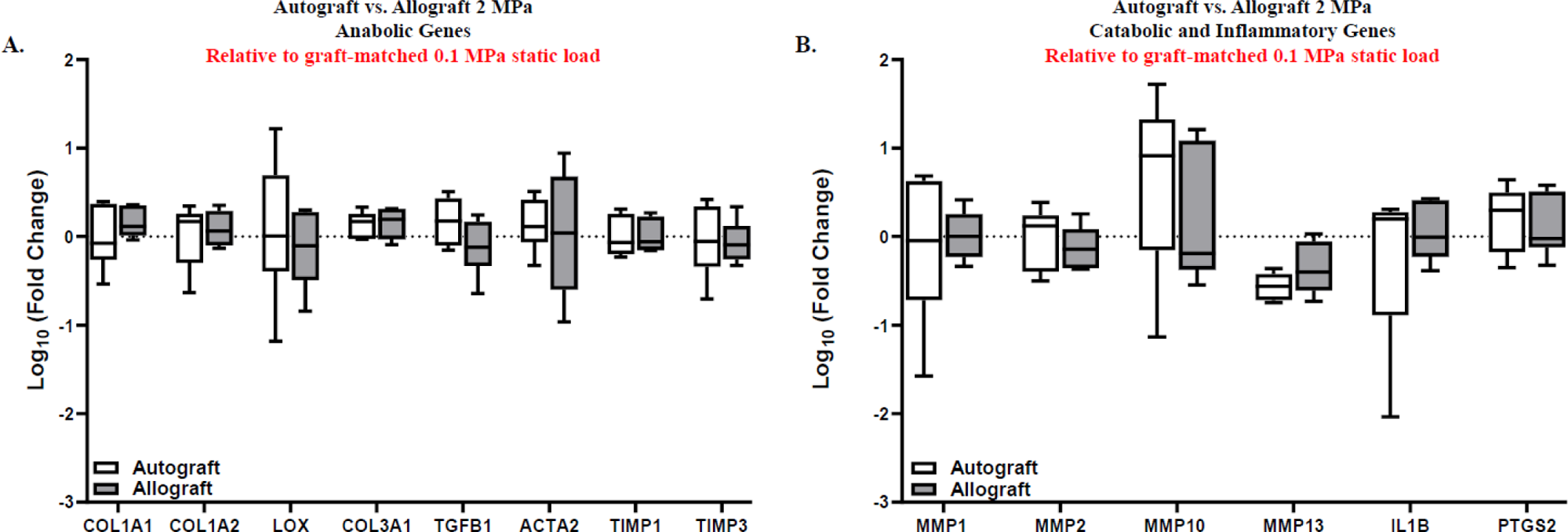
Comparison of autograft and allograft reconstructions gene expression to cyclic load. RT-qPCR analysis of autograft and allograft reconstructions cyclically loaded relative to their graft-matched static load. (A) Gene expression of anabolic markers. (B) Gene expression of catabolic and inflammatory markers. (n = 5-6 for cyclically loaded autografts and n = 6 for cyclically loaded allografts). Data represented as box and whiskers plot with the whiskers representing the min and max data.

Given the lack of gene expression changes in response to loading, we also directly compared the gene expression response of both autografts and allografts to data collected previously from native ACLs^36^. At baseline, we found that both graft types exhibited differential expression of multiple anabolic and catabolic genes compared to statically loaded native ACLs (**Supplementary Fig. S1**). Specifically, allografts upregulated multiple anabolic genes (*COL1A2, COL3A1*) and catabolic genes (*MMP2, MMP13*) compared to the statically loaded native ACL (p < 0.05) with a trending downregulation of the anabolic gene *ACTA2* (p < 0.10). Autografts also upregulated catabolic genes (*MMP2, MMP13*) (p < 0.05) with the anabolic gene *COL1A1* trending toward upregulation (p < 0.10). With cyclic loading to 2 MPa, autografts downregulated the expression of the catabolic gene *MMP13* compared to the native ACL (p < 0.05) with the inflammatory gene *IL-1β* trending toward downregulation (p < 0.10) (**Supplementary Fig. S2**). For cyclically loaded allografts, the expression of the anabolic gene *TGFβ1* was significantly downregulated compared to the native ACL (p < 0.05).

Finally, to assess whether potential mechanosensitive gene expression changes in reconstruction were obfuscated by increased noise (i.e., biological variability), we also compared the variability of the gene expression data of the statically loaded grafts to prior data on the native ACL^36^ (**Supplementary Table 2).** For statically loaded autografts, there was significantly increased variability in anabolic genes *LOX* and *TIMP3* compared to statically loaded native ACL (p < 0.05). For statically loaded allografts, there was only one anabolic gene (*LOX*) that trended towards increased variability compared to statically loaded native ACLs (p < 0.10).

### Immune cell gene expression of ACL reconstructions to cyclic load

In the statically loaded samples, there was no differential gene expression of any of the immune cell genes in allografts compared to autografts (**Fig. 5A**). Additionally, there was no change in the expression of any of the immune cell genes with mechanical loading in autografts (**Fig. 5B**) or allografts (**Fig. 5C**). Furthermore, direct comparison of the gene expression response to mechanical loading between autografts and allografts found no difference for any of the immune cell genes (**Fig. 5D**).

**Figure 5:**
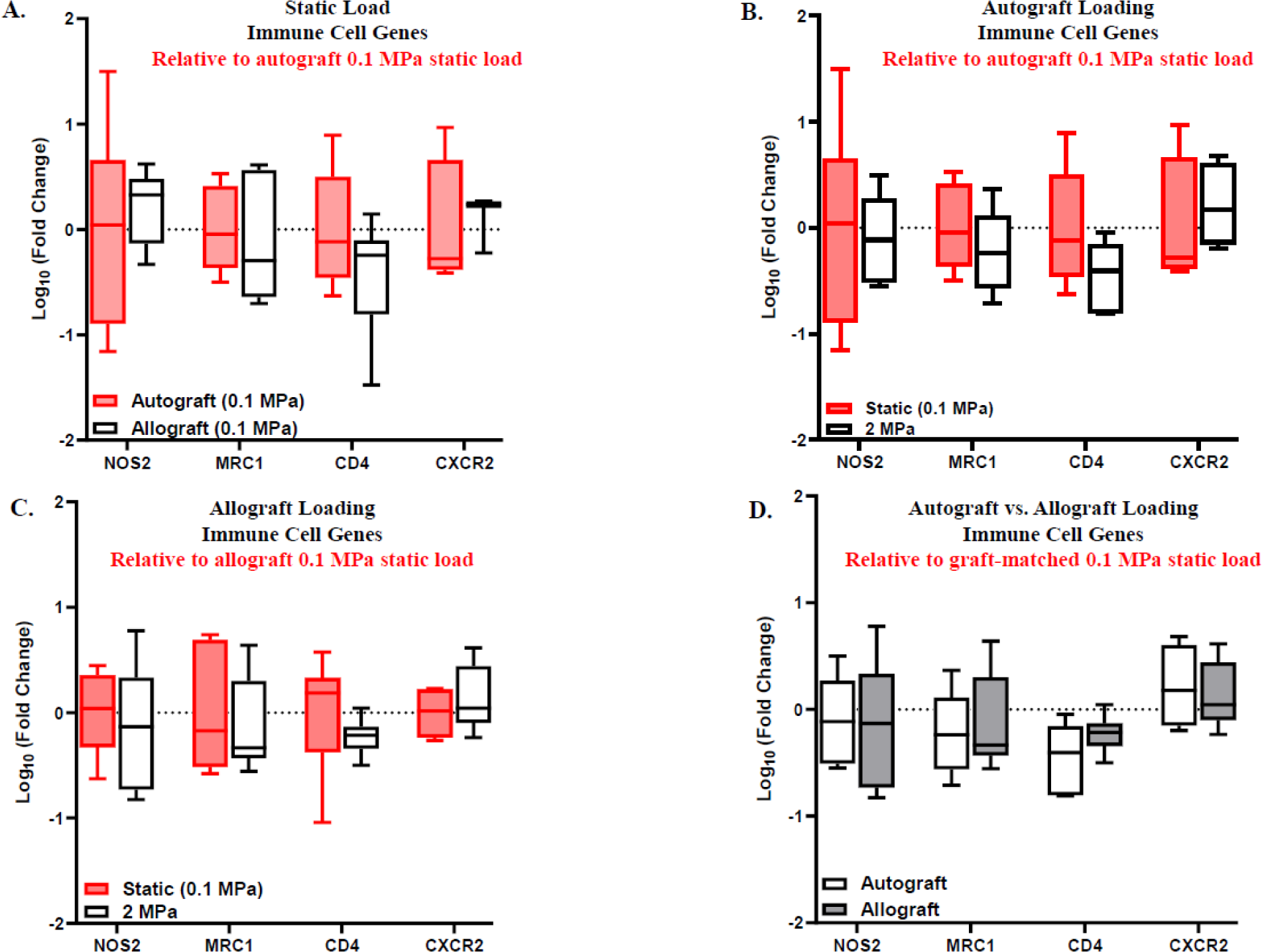
Comparison of immune cell gene composition between autografts and allografts. RT-qPCR analysis of immune cell gene composition between statically loaded autografts and allografts and cyclically loaded autografts and allografts. (A) Gene expression of statically loaded allografts relative to statically loaded autografts (represented by the red box and whisker plot). (B) Gene expression of autografts to cyclic load relative to statically loaded autografts (represented by the red box and whisker plot). (C) Gene expression of allografts to cyclic load relative to statically loaded allografts (represented by the red box and whisker plot). (D) Comparison of gene expression of statically loaded autografts and allografts relative to the graft-matched 0.1 MPa static load. (n = 4-6 for statically loaded allografts, n = 4-6 for cyclically loaded autografts, and n = 5-6 for cyclically loaded allografts). Data represented as box and whiskers plot with the whiskers representing the min and max data.

### Mechanical and physical comparison of reconstructions

The cross-sectional area and length of the allograft reconstructions (4.40 ± 1.7 mm^2^ and 8.34 ± 1.2 mm) were not significantly different compared to the autograft reconstructions (4.04 ± 2.0 mm^2^ and 8.17 ± 1.3 mm) (**Fig. 6A-B**). Additionally, there were no significant differences in the macroscale tissue strains between allograft (14.3 ± 7.4%) and autograft reconstructions (18.1 ± 4.7%) (**Fig. 6C**). Furthermore, there were no significant differences in the creep strain between allograft (3.4 ± 4.9%) and autograft (4.42 ± 3.7%) reconstructions (**Fig. 6D**).

**Figure 6:**
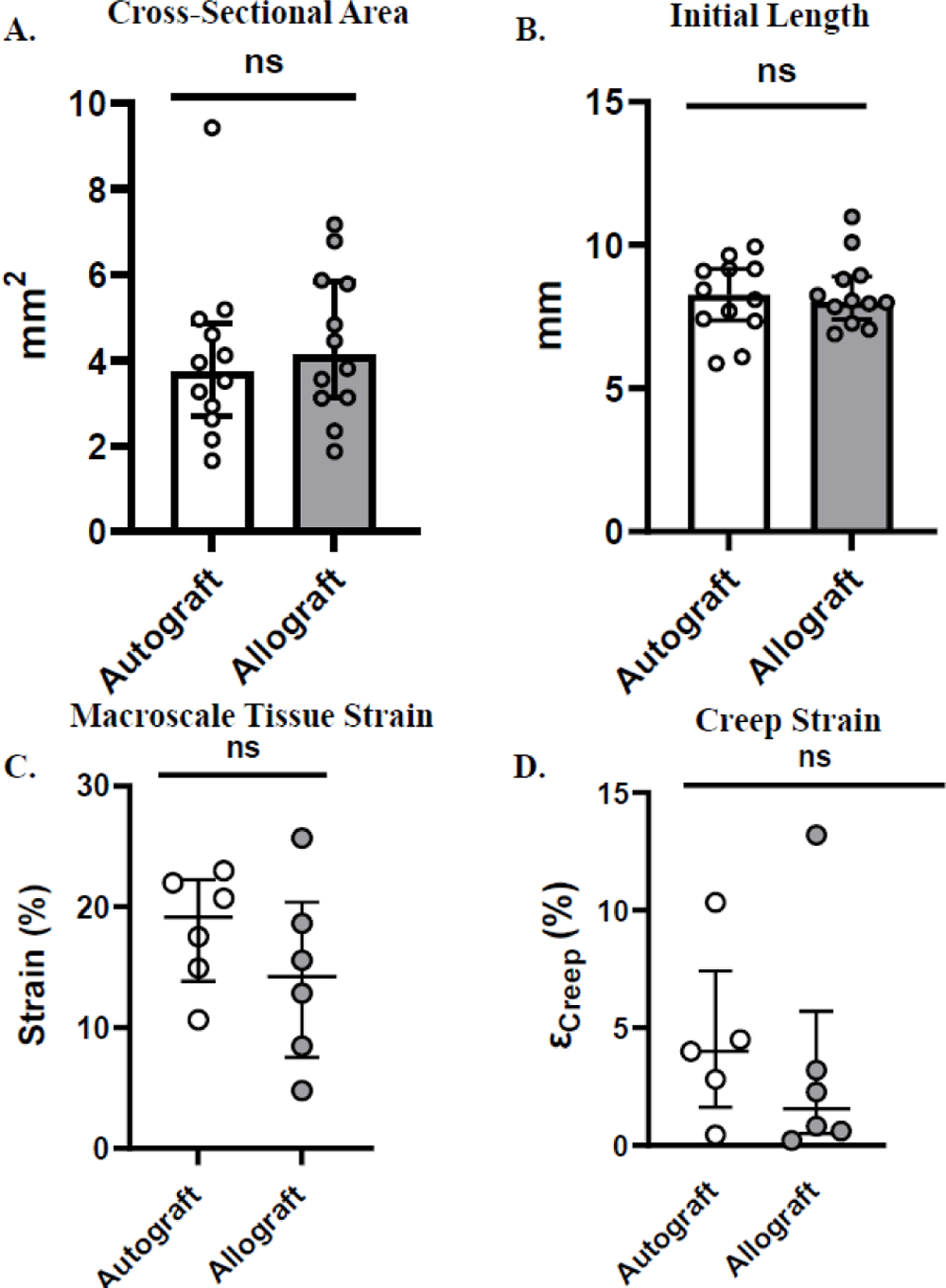
Physical and mechanical comparisons between autograft and allograft reconstructions. (A) Cross-sectional area of autograft and allograft reconstructions (n = 12). (B) Initial length of autograft and allograft reconstructions (n = 12). (C) Macroscale tissue strain of autograft and allograft reconstructions when cyclically loaded to 2 MPa (n = 5 for autografts and n = 6 for allografts). (D) Creep of autograft and allograft reconstructions when cyclically loaded to 2 MPa. Data represented by median and interquartile range.

## Discussion

The objective of this study was to compare the biological response of autograft and allograft ACL reconstructions to cyclic loading. We hypothesized that allografts would exhibit an impaired (i.e., more catabolic) remodeling response to load compared to autografts. Contrary to our hypothesis, we found that the remodeling response of allografts was not different than that of autografts in response to 2 MPa of cyclic loading. An important caveat is that a 2 MPa load may not be sufficient to induce a differential mechanobiological response between graft types. Indeed, there was no effect of loading on any gene for either graft type. However, 2 MPa was sufficient to demonstrate a difference in mechanobiology between native ACLs and ACL reconstructions (**Supplementary Fig. S2**). Furthermore, the macroscale tissue strains produced by 2 MPa in autografts and allografts were similar to the tissue strains in the native ACL at 4 MPa (**Supplementary Fig. S3A**), which did produce a mechanobiological response in prior work^36^. Importantly, the creep strain levels were similar between both graft types and native ACLs (**Supplementary Fig. S3B**), which suggests that there was not substantial slippage at the bone tunnels during the cyclic loading and that the measured macroscale strains reflect the actual strain in the tissue midsubstance. Finally, we did not load the reconstructions at a higher stress level because the grafts would fail at the bone interface, suggesting that 2 MPa was the max physiological load that the grafts could handle. Together, this suggests that allografts and autografts do not exhibit a differential response to loading early in the remodeling process and that deficient mechanobiology cannot explain the impaired remodeling of allograft reconstructions.

While no prior study has directly compared the mechanobiology of allografts and autografts, our findings are mildly consistent with the literature. For example, a prior comparison of the mechanical properties of ACL reconstructions in rabbits at 8-weeks found no differences in stiffness between graft types,^47^ which is consistent with the similarity in macroscale strains measured in our allograft and autograft reconstructions. However, they found that the maximum tensile load strength of the grafts was approximately 30 N, which is roughly equivalent to 7.5 MPa and significantly greater than what we could achieve with our grafts. This may be due to the fact that our samples were cyclically loaded while the samples in the prior study were tested under a single ramp to failure test. Additionally, we found that the baseline expression of remodeling genes was not different between graft types, which is not consistent with prior data^25^. Specifically, a previous study of gene expression in rabbit ACL reconstructions found that *MMP1*, *MMP2*, and *LOX* were upregulated in freshly harvested allografts compared to autografts^25^. However, the differences in expression of these genes between graft types was small compared to the differences between either graft and native ACLs, which is consistent with our statically loaded data (**Supplementary Fig. S1**). Therefore, our data is likely representative of the mechanobiology of rabbit ACL reconstructions.

A secondary finding in this work is that allografts did not exhibit markers of a heightened immune response compared to autografts. Specifically, we found no differences in any of our immune cell gene markers between graft types at baseline or in response to cyclic load (**Fig. 5**). While there is some conflicting data in the literature^24,31^, our findings are in agreement with several previous studies suggesting that allografts do not elicit a heightened immunological response^33,48,49,50,51,52,53^. Still, it is also possible that we did not see a differential immune cell markers between graft types because the immune cells may have already migrated out of the tissue by 8 weeks post-reconstruction. For example, previous work demonstrated that macrophage accumulation peaks in the early healing process and significantly decreases by 28 days after surgery^54^. Another reason we may not have seen a difference may be that we may not have chosen the right marker genes, and we did not perform more rigorous flow cytometry. Immune cell subtypes are extremely complicated and categorizing macrophages broadly as M1 and M2 is not sufficient. Specifically, a previous study identified a specific inflammatory macrophage population in ACL reconstructions^55^. Together, this demonstrates that more research is necessary to better understand the immunological differences between graft types.

A final limitation to this study was that there was a lot of variability in the reconstruction and joint quality sample-to-sample. Specifically, some animals exhibited a large amount of knee scarring and scar tissue formation specifically around the reconstruction. Furthermore, some of the reconstructions were frayed or exhibited multiple separate bundles. While these variations were not specific to graft type and occurred equally between autografts and allografts, they did make measuring the physical dimensions and proper loading of the reconstructions prone to inconsistency. Nevertheless, we saw minimal differences in variability in the statically loaded reconstructions compared to the statically loaded native ACL (**Supplementary Table 2**), which suggests that the variability of the surgeries did not correlate to increased gene expression variability.

In summary, our findings suggest that there is not a mechanobiological difference between autografts and allografts. Specifically, we did not find a differential expression of markers of tissue remodeling in response to cyclic loading between graft types. While these results do not support the overall hypothesis that the reason for impaired remodeling and increased rupture of allografts in active individuals is due to deficient mechanobiology, future work is needed to rule out this hypothesis. Specifically, reinforcement of the bone insertions to load the grafts at higher stress levels would help identify the effect of loading magnitude on these findings. Additionally, future work could load for longer durations to potentially induce a differential mechanobiological response between graft types. Finally, future work could better characterize the immune cell composition between graft types by obtaining samples at earlier time points or use more precise techniques like flow cytometry or single cell RNA-sequencing. Together, these data could provide insight into the reason for increased allograft reruptures and potentially offer approaches to improve allograft performance.

## Supporting information

Supplementary Figures and Tables with Legend

## Acknowledgements

We would like to thank the veterinary staff at Penn State for assisting in surgeries and taking care of the rabbits after surgery. We also would like to acknowledge the Genomics Core at Penn State for running the PCR plates.

