## Supplementary Figures and Tables with Legend for "Allograft and Autograft Anterior Cruciate Ligament Reconstructions Exhibit a Similar Biological Response to Cyclic Loading"

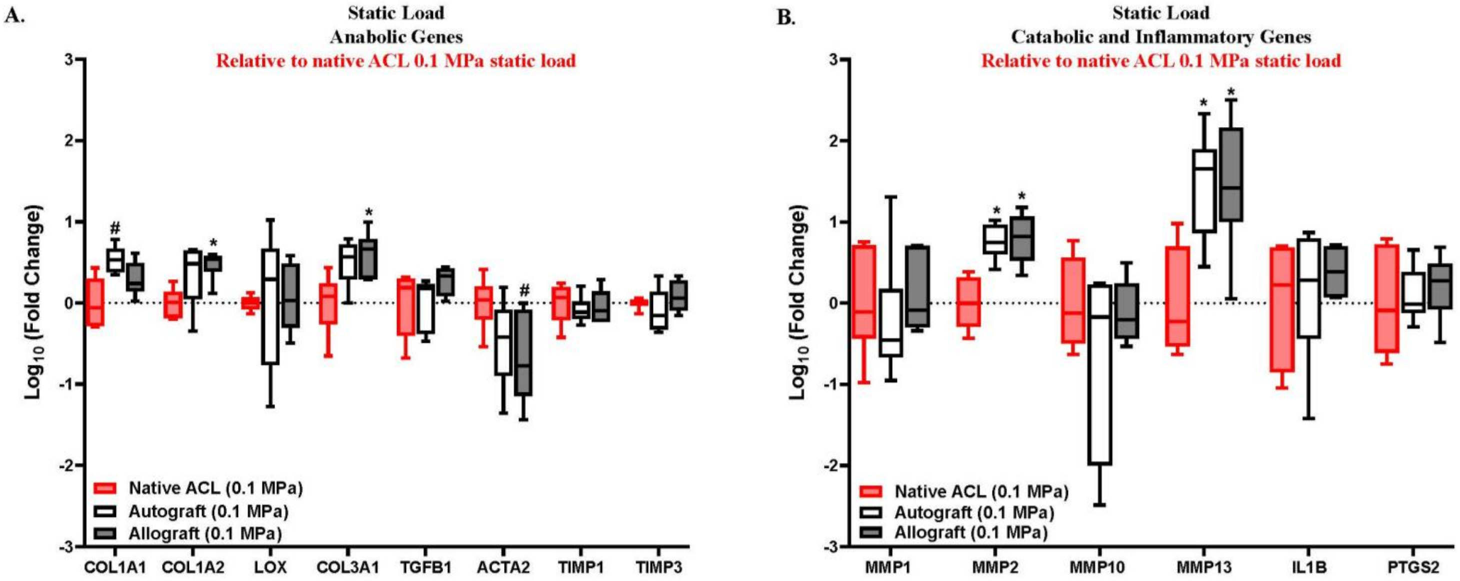


**Supplementary Figure 1:** Comparison of reconstructions and native ACL gene expression at baseline. RT-qPCR analysis of statically loaded autograft and allograft reconstructions relative to statically loaded native ACL (represented by the red box and whisker plots). (A) Gene expression of anabolic markers. (B) Gene expression of catabolic and inflammatory markers (n = 5-6 for autografts and n = 6 for allografts). *p < 0.05, #p < 0.10. Data represented as box and whiskers plot with the whiskers representing the min and max data. Data for native ACLs obtained from prior study[36].

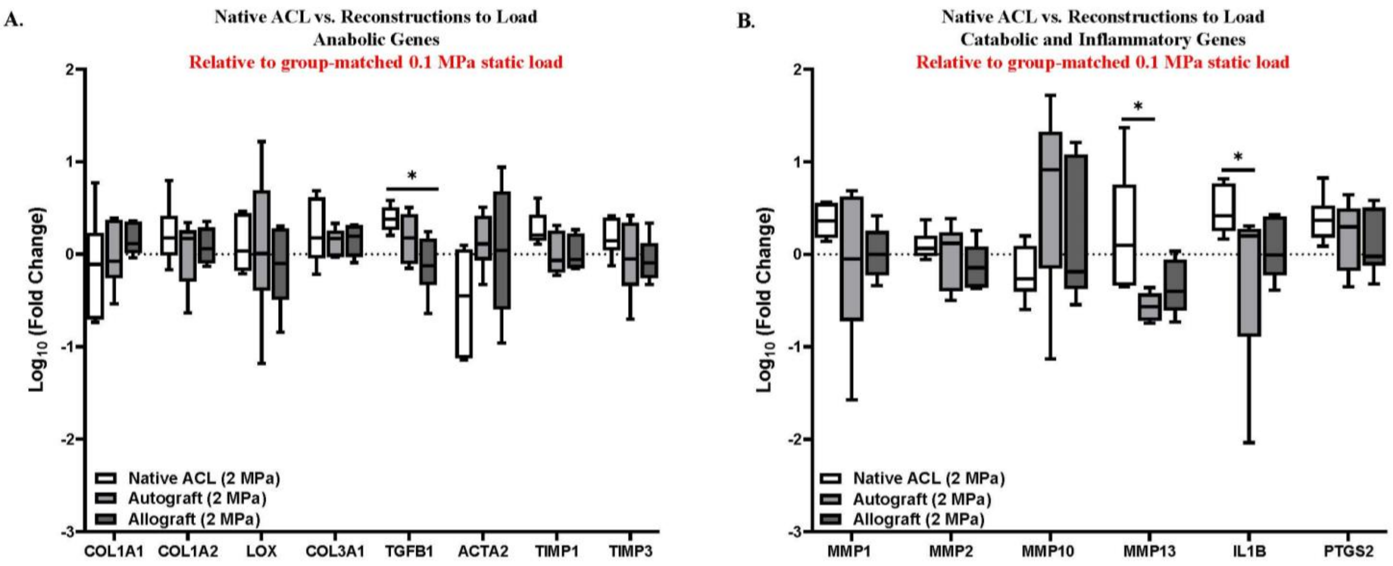


**Supplementary Figure 2:** Comparison of reconstructions and native ACL gene expression to cyclic load. RT-qPCR analysis of autograft and allograft reconstructions, and the native ACL cyclically loaded relative to their group-matched static load. (A) Gene expression of anabolic markers. (B) Gene expression of catabolic and inflammatory markers (n = 5-6 for autografts, n = 6 for allografts, and n = 6 native ACL). *$p < 0.05$. Data represented as box and whiskers plot with the whiskers representing the min and max data. Data for native ACLs obtained from prior study[36].

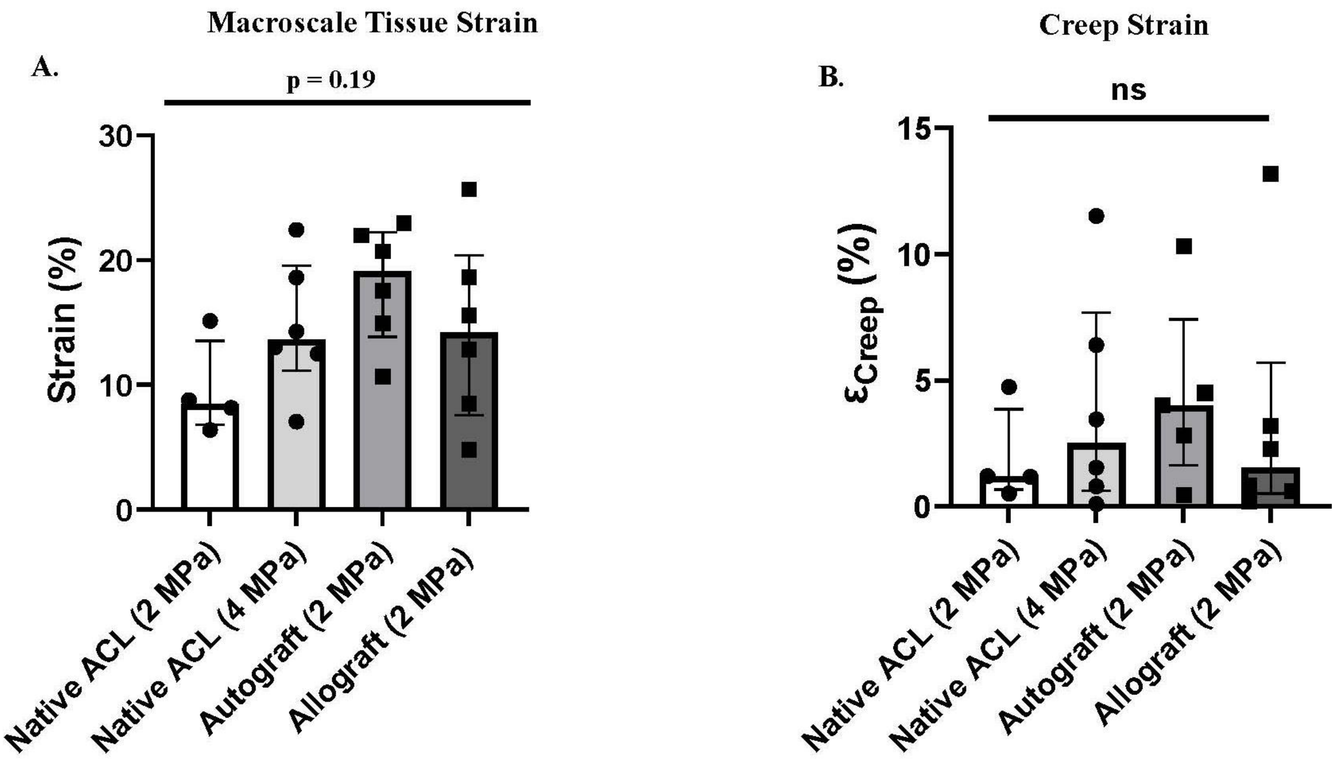


**Supplementary Figure 3:** Comparison of macroscale tissue strains between reconstructions and the native ACL. (A) Macroscale tissue strains and (B) creep strains of autografts and allografts cyclically loaded to 2 MPa, and the native ACL cyclically loaded to 2 and 4 MPa (n = 5 for autografts, n = 6 for allografts, n = 4 for 2 MPa native ACL, n = 6 for 4 MPa native ACL). Strains were compared between the reconstructions and the native ACL utilizing a Kruskal Wallis test. Data represented by median and interquartile range. Data for native ACLs obtained from prior study[36].

**Supplementary Table 1:** TaqMan probe gene ID and average efficiencies.

| Gene | Gene ID | Probe Efficiency |
|---|---|---|
| **COL1A1** | Oc03396073_g1 | 84% |
| **COL1A2** | Oc03396112_m1 | 78% |

| | | |
|---|---|---|
| **LOX** | Oc03398373_m1 | 85% |
| **COL3A1** | Oc03398373_m1 | 81% |
| **TGFB1** | Oc04176122_u1 | 81% |
| **ACTA2** | Oc03399251_m1 | 85% |
| **TIMP1** | Oc03397606_m1 | 79% |
| **TIMP3** | Oc04096947_m1 | 80% |
| **MMP1** | Oc04250657_m1 | 82% |
| **MMP2** | Oc03397553_m1 | 82% |
| **MMP10** | Oc03396490_m1 | 90% |
| **MMP13** | Oc03396899_m1 | 87% |
| **IL1B** | Oc03823250_s1 | 82% |
| **PTGS2** | Oc03398295_m1 | 81% |
| **NOS2** | Oc04096962_gH | 81% |
| **MRC1** | Oc06778204_g1 | 82% |
| **CD4** | Oc03823617_s1 | 83% |
| **CXCR2** | Oc04957973_m1 | 85% |
| **GAPDH** | Oc03823402_g1 | 78% |

**Supplementary Table 2:** p values from F-tests to compare variability of statically loaded

| Gene | Native ACL vs. Autograft | Native ACL vs. Allograft |
|---|---|---|
| COL1A1 | 0.301 | 0.714 |
| COL1A2 | 0.229 | 0.978 |
| LOX | 0.005* | 0.0983# |
| COL3A1 | 0.522 | 0.714 |
| TGFB1 | 0.576 | 0.233 |
| ACTA2 | 0.301 | 0.513 |
| TIMP1 | 0.419 | 0.855 |
| TIMP3 | 0.035* | 0.172 |
| MMP1 | 0.968 | 0.839 |
| MMP2 | 0.103 | 0.854 |
| MMP10 | 0.307 | 0.839 |
| MMP13 | 1.000 | 0.839 |
| IL1B | 0.870 | 0.104 |
| PTGS2 | 0.378 | 0.355 |
